# Gene expression profiles based flux balance model to predict the carbon source for *Bacillus subtilis*

**DOI:** 10.1101/842518

**Authors:** Kulwadee Thanamit, Franziska Hoerhold, Marcus Oswald, Rainer Koenig

## Abstract

Finding drug targets for antimicrobial treatment is a central focus in biomedical research. To discover new drug targets, we developed a method to identify which nutrients are essential for microorganisms. Using ^13^C labeled metabolites to infer metabolic fluxes is the most informative way to infer metabolic fluxes to date. However, the data can get difficult to acquire in complicated environments, for example, if the pathogen homes in host cells. Although data from gene expression profiling is less informative compared to metabolic tracer derived data, its generation is less laborious, and may still provide the relevant information. Besides this, metabolic fluxes have been successfully predicted by flux balance analysis (FBA). We developed an FBA based approach using the stoichiometric knowledge of the metabolic reactions of a cell combining them with expression profiles of the coding genes. We aimed to identify essential drug targets for specific nutritional uptakes of microorganisms. As a case study, we predicted each single carbon source out of a pool of eight different carbon sources for *B. subtilis* based on gene expression profiles. The models were in good agreement to models basing on ^13^C metabolic flux data of the same conditions. We could well predict every carbon source. Later, we applied successfully the model to unseen data from a study in which the carbon source was shifted from glucose to malate and *vice versa*. Technically, we present a new and fast method to reduce thermodynamically infeasible loops, which is a necessary preprocessing step for such model-building algorithms.

**SIGNIFICANCE:** Identifying metabolic fluxes using ^13^C labeled tracers is the most informative way to gain insight into metabolic fluxes. However, obtaining the data can be laborious and challenging in a complex environment. Though transcriptional data is an indirect mean to estimate the fluxes, it can help to identify this. Here, we developed a new method employing constraint-based modeling to predict metabolic fluxes embedding gene expression profiles in a linear regression model. As a case study, we used the data from *Bacillus subtilis* grown under different carbon sources. We could well predict the correct carbon source. Additionally, we established a novel and fast method to remove thermodynamically infeasible loops.

## INTRODUCTION

Infectious diseases are a severe burden of public health. They are caused by micro-organisms or viruses and responsible for premature death and disability. Typically, they are treated by a broad range of antimicrobials. However, pathogenic micro-organisms are challenging to treat and in particular pathogens which hide inside host cells. For example, osteomyelitis is an infection in the bone marrow and mainly caused by *Staphylococcus aureus* (*S. aureus*) ^(1-3)^. These bacteria express adhesins to promote adherence to osteoblasts or collagen and host inside the macrophages in the bone marrow. After an initial short duration of trying to get resistant, typically, the host cells get resilient and the pathogen enters a dormant state ^(4)^ or the pathogens colonize in biofilms ^(3, 5, 6)^. The infection is difficult to treat since antibiotics must penetrate through the host cell or biofilm to eradicate the bacteria. Hence, such treatments result in low success rates. Furthermore, antibiotic resistance contributes to worsening the situation ^(7, 8)^. So, new drug targets or interventions are needed. For living organisms, it is essential to get adequate nutrients maintaining their metabolism. This also applies to these pathogens even though in low metabolic rates when they are in such a dormant state.

Besides this, constraint-based modeling (CBM) or flux balance analysis (FBA) has been used to investigate metabolic networks. The fluxes of the reactions are derived from models basing on physiochemical constraints. Assuming a steady-state condition, these constraints are obtained by the stoichiometry of each reaction assuming mass balance inside the cell for each (inner) metabolite. Additional constraints may be derived from thermodynamic constraints implying the directionality and enzyme capacity estimating a maximal enzymatic rate (Vmax). Steady state bypasses the need for reaction kinetic parameters facilitating to construct metabolic models on a genome scale without the experimentally demanding determination of these parameters ^(9-11)^. FBA has been served as a tool to provide *in silico* simulations which allows researchers to discover new targets and support further studies. It has been widely applied to model metabolic fluxes in a scalable way, particularly, in the field of metabolic engineering or agriculture to enhance the yield of targeted products ^(12-15)^. FBA has also been applied in health sciences investigating pathomechanisms of diseases to find new drug targets ^(16-19)^. In FBA models, a typical objective function of microorganisms or tumor cells is to maximize biomass production. However, we often see that utilizing only stoichiometric data of cell reactions is inadequate to achieve good phenotype predictions for a specific condition. Therefore, experimental data, for example, metabolite concentrations derived from ^13^C labeling experiments or transcriptomic profiles, have been integrated to improve flux estimations.

Metabolic fluxes can be derived from employing ^13^C-labeled tracers. Although such a ^13^C flux analysis is powerful, the experimental implementation can be laborious and difficult ^(20)^. Moreover, in a complex environment such as osteomyelitis, it is experimentally challenging to identify which metabolite belongs to the host or pathogen, particularly, when the bacteria are in a dormant state. Therefore, transcriptional regulation has been in the spotlight for computational biologists to utilize this information, as this experimental data is much easier to obtain than metabolic flux data from ^13^C isotopic tracer analyses ^(21-23)^; also, in clinical settings where ^13^C labeling experiments are hard to achieve. We followed the approach employing FBA prediction by integrating gene expression profiles to identify the nutritional needs of a bacteria based on gene expression data. As a case study, we demonstrated our concept using publicly available gene expression data from the Gram-positive, competent and well-studied bacterium *Bacillus subtilis* (*B. subtilis*) under growth conditions with different carbon sources ^(24, 25)^. We aimed to predict the correct carbon sources basing on the transcriptional profiles at these specific conditions.

## MATERIALS AND METHODS

### 1. Data assembly

#### 1.1. The experimental data of the first study (first dataset)

Published microarray gene expression data of *B. subtilis* strain BSB1, was used. BSB1 is a derivative of strain 168. The data was taken from a supplementary Table (Table S2) of the original publication ^(25)^. The data based on tilling arrays covering the whole genome of *B. subtilis* 168 ^(25)^. *B. subtilis* was grown in minimal medium in eight different carbon source conditions (glucose, fructose, gluconate, glutamate/succinate, glycerol, malate, malate/glucose, pyruvate) ^(25)^. In addition, we used metabolic flux data basing on ^13^C isotope labeling experiments by Chubukov *et al* of the same eight carbon source conditions ^(26)^. The data was taken from Supplementary Table S4 of the publication. This data will also be denoted as the first dataset in the following.

#### 1.2. Experimental data of the second study (second dataset)

To validate our model with a separate, unknown dataset, we used publicly available gene expression data and ^13^C tracer based metabolic flux data from a time-series of two nutritional shifts, i.e. from glucose to glucose plus malate, and from malate to glucose plus malate ^(24)^. This data will also be denoted as the second dataset in the following. Gene expression data and ^13^C metabolic flux data were generated using the same techniques as described above. *B. subtilis* was grown in minimal medium on a single carbon substrate until an OD600 of 0.5 was achieved. Then, the other substrate (glucose or malate) was added to the culture to assess the bacterial behavior 0 (before the addition of the other substrate), 5, 10, 15, 25, 45, 60 and 90 minutes after the nutritional shift. Both gene expression and ^13^C metabolic flux data were taken from the BaSysBio database (https://basysbio.ethz.ch/openbis/basysbio_openbis.html).

### 2. Data pre-processing

We used the gene expression data of the first and second dataset. It had been preprocessed by computing the median of the estimated transcription signal of the probes assigned to the according gene ^(24, 25)^. The gene expression data in the second dataset had been further quantile normalized ^(24)^. All gene expression levels were provided in log2 transformation ^(24, 25)^. To obtain the gene symbols, BSU numbers were matched with gene symbols using bioDBnet version 2.1 ^(27)^. In the first dataset, each condition contained three biological replicates and we used all of them. For the majority of the time points of the second dataset, three biological replicates were generated while some time points contained two biological replicates. For each condition/time point, gene expression levels from the available replicates were averaged in our study. To map gene expression values to proteins and reactions, we used the gene-protein-reaction (GPR) mapping from the original publication of Chubukov *et al* ^(26)^ and the metabolic network of *B. Subtilis* 168 available from the BiGG Models database (BiGG ID iYO844) ^(28)^. We compared the mappings with the information from Uniprot ^(29)^ and KEGG ^(30-32)^ and corrected it if stated otherwise in these databases. Additionally, we found literature about two more genes (lrgA, lrgB) coding for a pyruvate transporter and added them to the corresponding reactions in the GPR mapping ^(33)^. The GPR mapping we used is provided in Table S1 in the Supplementary Information. ^13^C metabolic flux data from Chubukov *et al* ^(26)^ and Buescher *et al* ^(24)^ were used as published, without further processing.

### 3. Model building

#### 3.1. Building the metabolic model

To work with our mixed-integer linear programming/constraint-based model implementation, we transferred the iYO844 model of *B. subtilis* from Matlab to R (stoichiometric matrix, lower and upper bounds, reversibility, metabolite, and reaction names). Initial trials showed that we could not find any solution in the solution space when we tried to fit flux values from ^13^C metabolic flux data when allowing only one exchange reaction flux to be non-zero, i.e. from the specific transporter of the according carbon source. However, it was possible to find a feasible solution when we allowed fluxes from other exchange reactions to enter the system. Although the solution was found, the flux values from other exchange reactions were substantially high, which was unrealistic. Hence, we set a reasonable boundary for these exchange reactions letting the solution deviate from ^13^C metabolic flux data by maximal 0.1. This led to the lowest possible sum of fluxes of 0.688 which was applied to restrict these exchange reaction fluxes entering the system. The list of all exchange reactions besides designated carbon sources is provided in Table S3 in the Supplementary Information.

#### 3.2. Defining the set of reactions for the optimization criterion

As described below, we benchmarked our models with well-defined gold standards, i.e. the flux values based on the ^13^C labeling data from the original publications. This gold standard data was available for 40 reactions mostly covering central energy metabolism. Hence, these reactions were used for the optimization of our metabolic model explained in the next section. These reactions are called core reactions in the following. To improve the model predictions, we added a selection of further reactions to the optimization function of our model, called associated reactions in the following. We wanted to add associated reactions following three criteria, (1) they needed to be reactions which were directly connected to the core reactions in central energy metabolism or amino acid biosynthesis, (2) important metabolites in glycolysis or tricarboxylic acid (TCA) cycle (e.g. glyceraldehyde 3-phosphate, pyruvate, oxaloacetate, α-ketoglutarate) should participate in these reactions, and (3) at least one of the associated genes to the reactions needed to be differentially expressed in at least one out of the eight carbon sources of the first dataset when compared to the expression of *B. subtilis* in the control medium (*B. subtilis* grown in LB medium) ^(24)^. For this, T-tests were performed comparing the expression values of the according gene in each specific carbon source condition *versus* its expression in samples from an LB medium (control). The Benjamini-Hochberg method was used to correct for multiple testing across all genes ^(34)^. The p-value cutoff was 0.05. By this, we assembled 119 genes and 138 reactions in total (Table S1).

### 4. Formulating the optimization criterion

Assuming that the metabolic flux correlates linearly with the expression of the gene coding for the responsible enzyme of the according reaction, we linearly mapped gene expression values to predicted fluxes, formulated within the following optimization problem.

Let 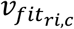 represent a gene expression-based flux for reaction *ri* in condition *c*. 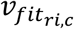 is derived by using information from gene expression data and the flux range,

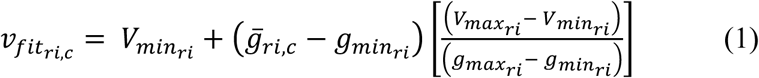

where 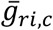 is the gene expression value of the gene associated to reaction *ri* in condition *c*, 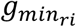 is the minimum gene expression value and 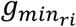 the maximum gene expression value across all conditions. 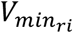 is the minimum and 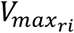 the maximum possible flux across all conditions, obtained from the gold standard (^13^C metabolic flux) for the core reactions (CR). For the associated reactions (AR), the minimal and maximal flux values are derived using flux variability analysis (FVA) explained below.

Under the FBA framework, we assumed that the metabolic network is in steady state; there is no accumulation of mass, which means no change of metabolite concentration over time. While *S*_*r*_ is the stoichiometric matrix of the metabolic network, *v*_*r,c*_ represents the predicted flux for reaction *r* in condition *c* in the metabolic network. The variable *v*_*r,c*_ must satisfy the constraints from the stoichiometry, as well as lower *lb*_*r*_ and upper bounds *ub*_*r*_, i.e.

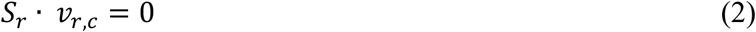

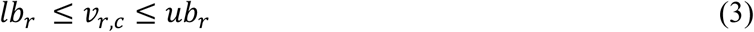

Subject to constraints (1) to (3), we formulated the optimization problem by

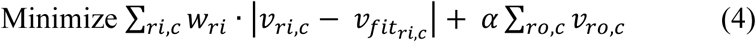

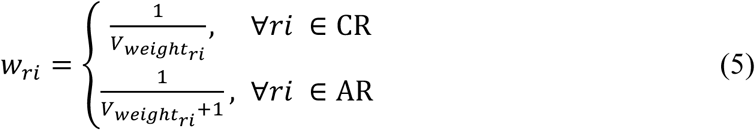

The formulated objective function is a trade-off between two optimization criteria. The first term, 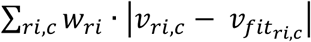, minimizes the error between the predicted flux *v*_*ri,c*_ and gene expression-based flux prediction 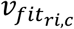. The weight *w*_*ri*_ is introduced to adjust the term through equation (5). By applying *w*_*ri*_, we did not need to consider AND/OR rules when a reaction *ri* was associated with more than one gene. The predicted flux *v*_*ri,c*_ was adjusted by averaging the gene expression values using the weight *w*_*ri*_ for each gene encoding this reaction *ri*. 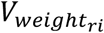 was obtained by selecting the maximum of absolute values of maximum or minimum flux from ^13^C metabolic flux (CR) or FVA calculated flux (AR). The weight was set as the reciprocal of this value to equal out the nominator in equation (3) making reactions with small and high variances of fluxes equally important to the objective function. The associated reactions were down-weighted by adding the constant +1 in the denominator. Moreover, we discarded reactions which 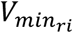 and 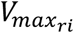 were zero. This resulted in lower numbers of reactions leading to 98 reactions basing on 116 genes (listed in Table S2).

The second term in formula (4), *α* Σ_*ro,c*_ *v*_*ro,c*_, aims to minimize the sum of all predicted fluxes *v*_*ro,c*_ from reactions being not CR nor AR coping for the problem of obtaining thermodynamically infeasible loops. The value of *α* was set to 0.01 and selected since the sum of flux *v*_*ro,c*_ was greatly reduced while the total model mapping error was not raised. The results of varying *α* values are provided in Fig.S1 in the Supplementary Information.

Notably, a biomass constraint was set for each different *c* based on publications of Chubukov *et al* and Buescher *et al* ^(24, 26)^. In our implementation, we opened the lower bounds for all eight carbon source exchange reactions at once to allow fluxes to enter the system due to the predictions based on the according gene expression profiles. For each carbon source, we took the maximum substrate rate reported in Chubukov *et al* ^(26)^ across all conditions. We set the lower (negative) bounds for all eight carbon source exchange reactions according to these values.

#### 4.1 Reducing the search space iteratively employing flux variability analysis (FVA)

To correctly map the expression data to the metabolic fluxes, we needed a realistic estimate of the lower and upper bounds for the reactions in the model. At different steady-state conditions, feasible minimum and maximum fluxes can differ from their initially set lower and upper bounds for each reaction in the metabolic network. FVA is a well-known technique to determine the maximal flux ranges ^(35)^. We applied FVA to the associated reactions to determine the minimum and maximum possible fluxes as follows. For each reaction *r* in condition *c*, in FVA, it is assumed that the metabolic network is in steady state and the stoichiometry is fulfilled, as referred to equation (2) and (3). FVA minimizes and maximizes the flux *v*_*r,c*_ to find an upper and lower bound for the respective reaction satisfying the FBA constraints (formula (2) and (3)).

In general, doing this for every reaction should narrow down the flux range of each reaction. However, we observed that the boundaries did not differ substantially after performing FVA for every associated reaction. To further reduce the solution space and limit its flexibility, we developed an iterative approach. The approach was part of the training scheme to narrow the flux ranges for the associated reactions, hence, we applied this only to the training data. The method works as follows:

a. 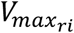 and 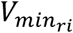 are acquired by the above-described FVA for each associated reaction.
b. Absolute values of 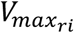 and 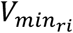 from each associated reaction are compared and the maximum of these values used as the representative maximal bound for this reaction. Representative maximal bounds from all associated reactions are used to rank the associated reactions. The reaction with the highest value is placed at the top position.
c. The first reaction *ri*, with i = 1 is selected.
d. 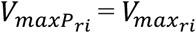 and 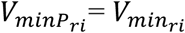 is set.
e. 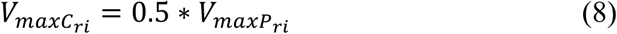

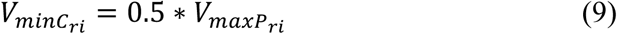

in which 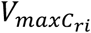 is the new maximal possible flux for the current iteration, 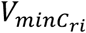 is the new minimal possible flux for the current iteration. 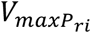 and 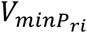 are reduced by half every iteration. Since the reaction can be unidirectional or bi-directional, 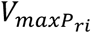 and 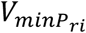 can have similar or different signs. Equation (8) and (9) are applied to reduce 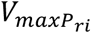 and 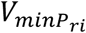. If 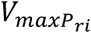 and 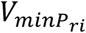 have the same sign, either equation (8) or (9) is used depending on the sign.
f. 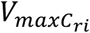 and 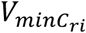 are applied as 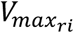 and 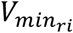 in formula (1) to mapping gene expression values to fluxes.
g. After optimizing the objective function in formula (4), the total model mapping errors are compared between the previous and the new iteration. If the total model mapping error from the previous run is greater, the algorithm proceeds with the next iteration and goes to step e).
h. The inner iterative process terminates for reaction i. The next reaction in the list is selected by setting i = i+1, and the algorithm proceeds with step d).
i. The algorithm terminates if the total model mapping error becomes stable or all reactions are processed.

With this approach, the set of 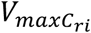 and 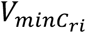 was chosen considering two criteria: 1) the total model mapping error from this iteration was lower compared to the previous iteration and 2) the predicted results of the fluxes for GapA and GapB reflected their expected activities when a glycolytic (glucose, fructose, gluconate, glycerol) or glucogenic (glutamate, succinate, malate, pyruvate) carbon source was taken up in the system. To be noted, in this study, we terminated the process when the total model mapping error did not decrease anymore and the predicted results of the fluxes for GapA and GapB did not improve (Table S4). This substantially reduced the computational time by 25%.

### 5. Reducing the number of thermodynamically infeasible loops (RED-TIL)

In constraint-based modeling, the thermodynamic loop law is often not taken into account to reduce computational complexity. The loop law is similar to Kirchhoff’s second law for electrical circuits ^(36)^. It states that at steady state there must not be any closed cycle or loop in the metabolic network with a non-zero net flux. Such loops would disregard the second law of thermodynamics and are hence thermodynamically infeasible. However, when optimizing the metabolic fluxes best matching to transcriptional profiles of the enzyme coding genes, such thermodynamically infeasible loops (TILs) may be exploited by during optimization to fit the fluxes to the expression values of the reactions. The problem of avoiding thermodynamically infeasible loops can be solved by imposing thermodynamic constraints such as standard-state free energy of reaction into the optimization. However, it is very challenging to acquire information for the whole metabolic network.

To solve this problem within the FBA framework, Schellenberger *et al* ^(37)^, introduced a method called loopless-COBRA (ll-COBRA). Il-COBRA removes TILs from the network. The method does not require additional thermodynamic information. It utilizes directions of flux distributions, which are readily available in every metabolic network. The direction of flux is then referred to a driving force of a reaction. If the direction is positive, the driving force of the reaction is negative and *vice versa*. Considering internal reactions, a loopless solution exists when the driving forces for all internal reactions that participate in any cycle are added up to zero. With this, the method finds the solution with no loop while generating a simpler mixed-integer linear programming (MILP) problem. Although the problem becomes less complex, it is still quite CPU intensive. We solved the TIL problem by a novel iterative procedure to speed up the process. The method is called RED-TIL in the following. After obtaining flux prediction results from the mapping procedure (see 4. Formulating the optimization criterion), the results were used as an input for a MILP problem to identify TILs and exclude them.

First, external reactions are not regarded. Applying a maximal flux value threshold (threshold = 0.01) for TILs to be allowed, the set of reactions *supp*(*v*) known as the support of *v* is assembled. *supp*(*v*) contains a subset of the internal reactions (*v* ≥ 0.01). We applied the value of 0.01 as a trade-off between CPU time and reasonable results. Next, an optimization problem is put up to determine the length of a minimum-containing TIL in the solution by

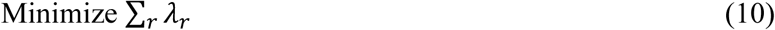

subject to

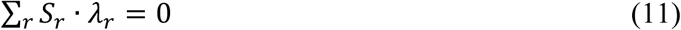

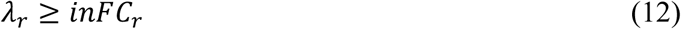

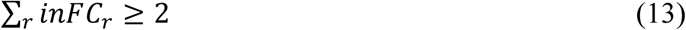

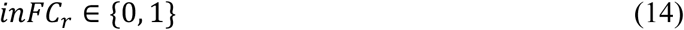

where *λ*_*r*_ is the flux of reaction *r* (∀*r* ∈ *supp*(*v*)), *S*_*r*_ is a stoichiometric matrix of the metabolic network with metabolites and reactions, *inFC*_*r*_ is a binary variable which equals to 1 for a reaction which is involved in the potential TIL. In a system that contains a TIL, there must be at least two reactions involved enforced by equation (13). If a solution of the problem put up by equation (10) – (14) is found, a TIL (of length k) is detected. A constraint is added not allowing this TIL by

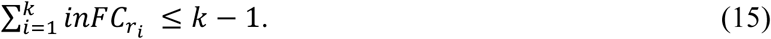

Equation (15) forces the algorithm to search for a solution that puts at least one of these variables 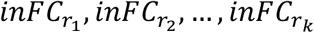 to 0 which leads to the TIL to be discarded from the solution. In the next optimization iteration, the mapping procedure is re-optimized using equation (1) to (5) together with the newly added constraint from equation (15), followed by finding new TILs employing the MIP problem described by equation (10) – (14). The algorithm stops when no TIL above the threshold can be found.

### 6. Validation of the model

The overview of the entire process is illustrated in Fig.1. We started by learning the model based on gene expression data as explained in the previous section obtaining the best parameter setting. Then, we predicted the primary carbon source for each of the eight carbon source conditions by selecting the transporter (one out of eight potential transporters) with the highest flux in the according condition. The prediction was validated by comparison to the known carbon source. In addition, we compared the flux predictions of the 40 core reactions with the fluxes of the original publication derived by ^13^C tracer analysis and quantified the similarity employing Pearson’s correlation. Furthermore, the model was applied to an unknown dataset, i.e. time series on spiked glucose on malate and spiked malate on glucose medium as described above.

**Figure 1.**
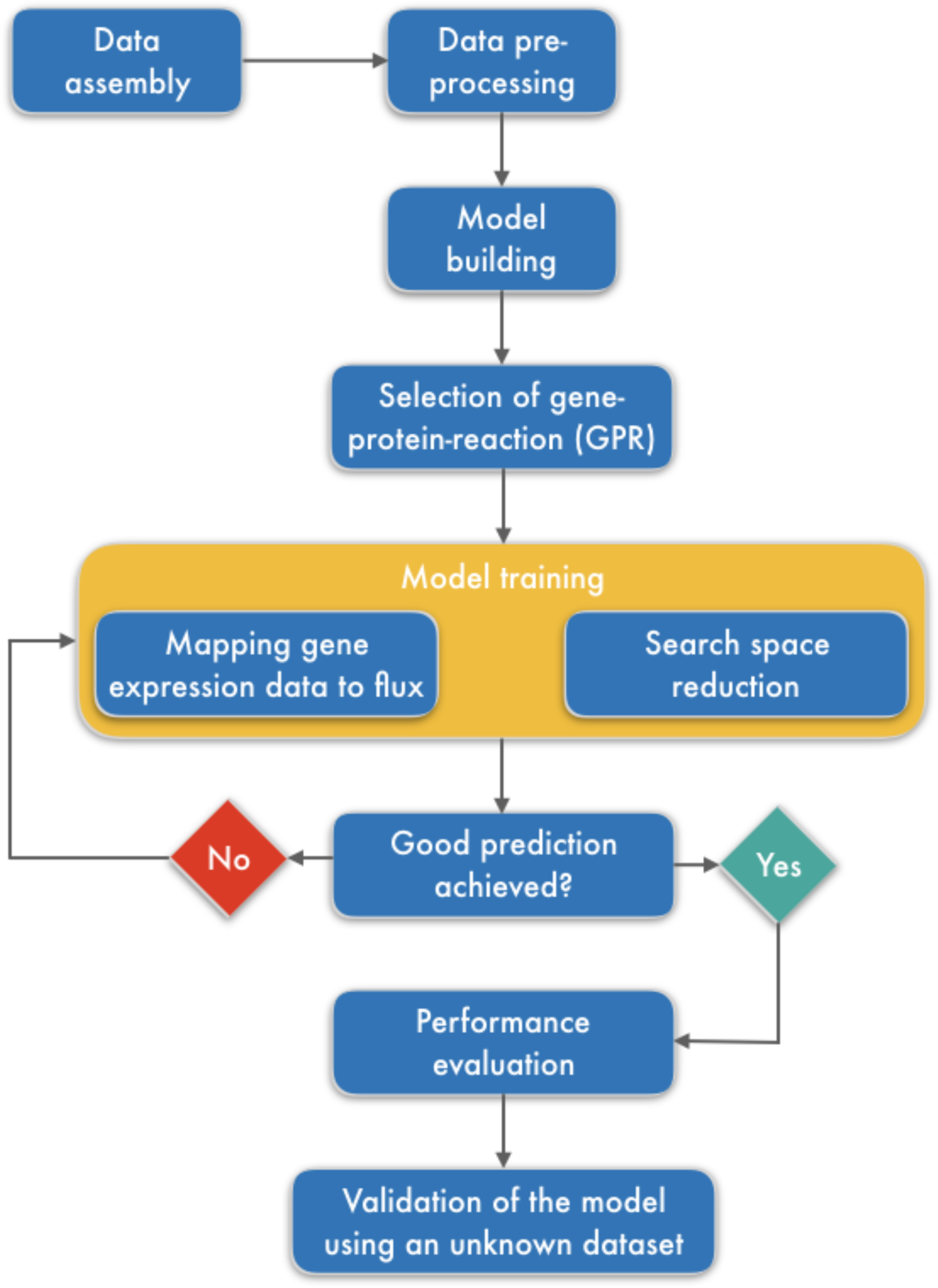
The workflow (see text).

### 7. Implementation

All analyses were performed using R version 3.3.3 (www.r-project.org). The Cobra toolbox version 2.0.6 and Matlab version R2013b (www.mathworks.com) were used to obtain the initial stoichiometric matrix, lower and upper bounds, reversibility information, and metabolite and reaction names from the metabolic network. All further analysis was performed using R. The Gurobi optimizer (version 7.0.1, www.gurobi.com) was used to solve all mixed-integer linear optimization problems.

## RESULTS

### Search space reduction improved flux predictions

To ensure that we considered only flux solutions within a thermodynamically feasible subspace, we trained the metabolic network model of *B. subtilis* by implementing the mapping approach together with the loopless RED-TIL method. Besides the core reactions, for which ^13^C metabolic flux data was available, we considered associated reactions for our gene expression based flux optimization. The associated reactions were neighbors of the core reactions that would be important for the carbon source prediction (see Materials and Methods for details). For these associated reactions, we performed FVA to reduce the maximal and minimal possible flux bounds. However, we observed that for many reactions, the resulting flux ranges did not substantially differ from the originally set upper and lower bounds. These high flux ranges were expected to not reflect realistic situations and needed to be further reduced. Hence, we developed a new method to reduce the fluxes iteratively by comparing the discrepancy between the fluxes derived from the expression values and the feasible fluxes from FBA, denoted as “total model mapping error” in the following. This led to new, considerably reduced bounds (Table S5). Notably, the total model mapping error decreased dramatically down to 4.6% of the original error (Fig.S2). We also compared the results from the model to the gold standard, i.e. the fluxes derived from ^13^C labeling experiments of Chubukov *et al* ^(26)^. We observed that after applying the iterative new approach for search space reduction, the prediction results improved significantly (p=0.0045, from Pearson’s correlation r = 0.57, σ = 0.28 to r = 0.65, σ = 0.27). The flux predictions before and after employing the search space reduction are provided in Table S6 and Table S7. Correlation values of before and after applying the search space reduction are listed in Table S8. Overall, the results show that employing our search space reduction method decreased the flux ranges considerably and led to more realistic flux ranges resulting in better flux predictions.

### Reducing thermodynamically infeasible loops (RED-TIL) employing an iterative novel method

Although ll-COBRA is a well-known method to efficiently remove TILs in constraints-based modeling, the main drawback is the computation time. ll-COBRA generates one large MILP problem and searches for an optimal solution within the thermodynamically feasible (loopless) region ^(37)^. We followed a novel method iteratively removing TILs to solve the same problem requiring less running time (RED-TIL). RED-TIL solves an FBA problem first and then uses this optimal solution as an input to identify and remove a TIL from the solution space, and the FBA problem is optimized again. The process is repeated until no TILs above a certain threshold (same as for ll-COBRA) is detected. To compare these approaches, we implemented both methods under the same R programming environment using the same numerical solver. Because of different approaches, it is possible that the outputs from RED-TIL (Table S7) and ll-COBRA (Table S9) were not exactly identical. Still, they were well comparable with respect to the error to the gold standard (MAERED-TIL = 1.66, MAEll-COBRA = 1.80). Additionally, the predicted fluxes from both methods in all eight conditions were generally similar (Fig.S3), resulting in high Pearson’s correlation (average r = 0.99). The exact Pearson’s correlation coefficients from all conditions are listed in Table S10. In turn, we observed striking differences in the running times between RED-TIL and ll-COBRA. We compared the running times and running time ratios between RED-TIL and ll-COBRA (Table 1). In general, the average runtime of RED-TIL was three times lower than ll-COBRA. It should be noted that speed differed under different conditions. As shown in Table 1, the running time ratios between RED-TIL and ll-COBRA varied from two to six times. Altogether, the results demonstrated that RED-TIL provided comparable outputs to ll-COBRA while consumed less runtime. It can be used as the alternative feasible method to remove TILs quicker.

**Table 1.** Running time comparison between ll-COBRA and RED-TIL

| Condition | II-COBRA<sup>a</sup> | RED-TIL<sup>a</sup> | Ratio<br>(II-COBRA/RED-TIL) |
| --- | --- | --- | --- |
| Glucose | 31.041 | 9.876 | 3.143 |
| Fructose | 29.7 | 8.736 | 3.4 |
| Gluconate | 29.447 | 12.392 | 2.376 |
| Glutamate/Succinate | 38.736 | 14.628 | 2.648 |
| Glycerol | 30.571 | 8.538 | 3.581 |
| Malate | 36.664 | 14.713 | 2.492 |
| Malate/Glucose | 28.163 | 4.783 | 5.888 |
| Pyruvate | 30.928 | 12.177 | 2.54 |
| **Average** | **31.906** | **10.73** | **2.974** |

<sup>a</sup> Running time in seconds on a workstation with Intel® Xeon CPU E5-2630 v4 and 64 GB
Ram

### Gene expression profile based flux balance model enables identifying the carbon source for *B. subtilis*

To predict the uptake of the major carbon sources, we restricted our model to central energy metabolism (glycolysis, tricarboxylic acid cycle (TCA cycle), pentose phosphate pathway (PPP), amino acid biosynthesis) as these biochemical pathways are mainly involved in catabolizing the potential carbon sources. The metabolic network of *B. subtilis* was trained with gene expression profiles from eight conditions (carbon sources: glucose, fructose, gluconate, glutamate/succinate, glycerol, malate, malate/glucose, pyruvate) ^(25)^. As we aimed to predict the carbon sources based on the given gene expression profiles, we set the upper and lower bounds of all eight carbon source transporter reactions to the maximal and minimal values of all conditions. The amount of flux for each carbon source was estimated running our model based on gene expression levels of the genes coding for the observed reactions.

To assess our flux prediction results (Table S7), we compared the z-scores of the transporters across each condition (Fig.2). We predicted the primary (and secondary) carbon source by the highest (and second highest) z-score of its corresponding transporter in a certain condition (Table 2). In six out of six single carbon source conditions, the predictions were correct. In line, regarding the two carbon source conditions, the primary carbon source was predicted correctly. The secondary carbon source was predicted correctly for malate/glucose. For succinate/glutamate, the glutamate transporter was slightly below the glycerol transporter (Fig.2). In summary, the method could well predict the carbon sources based on gene expression profiles of the respective conditions.

**Figure 2.**
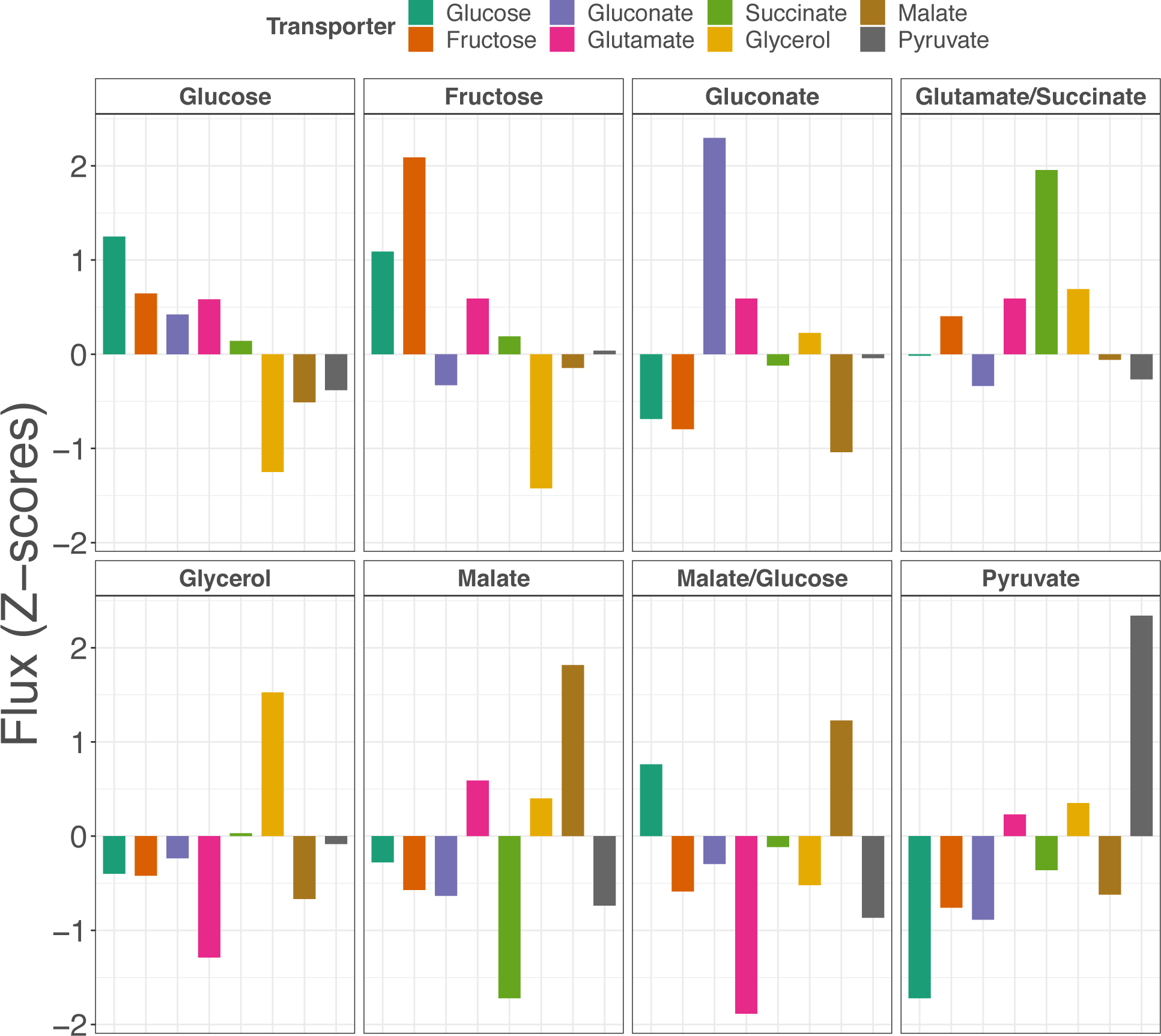
Predictions of the carbon source of the first dataset. A higher z-score indicates a higher probability for specific carbon source.

**Table 2.** Prediction of the carbon source for the first dataset

| Transporter | Condition |  |  |  |  |  |  |  |
| --- | --- | --- | --- | --- | --- | --- | --- | --- |
|  | Glucose | Fructose | Gluconate | Glutamate/Succinate | Glycerol | Malate | Malate/Glucose | Pyruvate |
| Glucose | 1 | 2 |  |  |  |  | 2 |  |
| Fructose | 2 | 1 |  |  |  |  |  |  |
| Gluconate |  |  | 1 |  |  |  |  |  |
| Glutamate |  |  | 2 | 3 |  | 2 |  |  |
| Succinate |  |  |  | 1 | 2 |  |  |  |
| Glycerol |  |  |  | 2 | 1 |  |  | 2 |
| Malate |  |  |  |  |  | 1 | 1 |  |
| Pyruvate |  |  |  |  |  |  |  | 1 |

### Gene expression profiles illustrate flux behavior inside the metabolic network

We compared our flux prediction results to the fluxes from the gold standard (derived by ^13^C labeling experiments from Chubokov *et al* ^(26)^). This comparison was performed for all 40 reactions, in which ^13^C metabolic flux data was available. For each of these reactions, the correlation of prediction and the gold standard values was calculated across all eight investigated carbon sources. As illustrated in Fig.3, most transporter reactions along with reactions in glycolysis and TCA cycle show good correlations with ^13^C metabolic flux data (r > 0.60), while reactions in PPP show lower correlations (r < 0.40). The average correlation across all 40 reactions is 0.65. Correlation coefficients of all 40 reactions are listed in Table S8.

**Figure 3.**
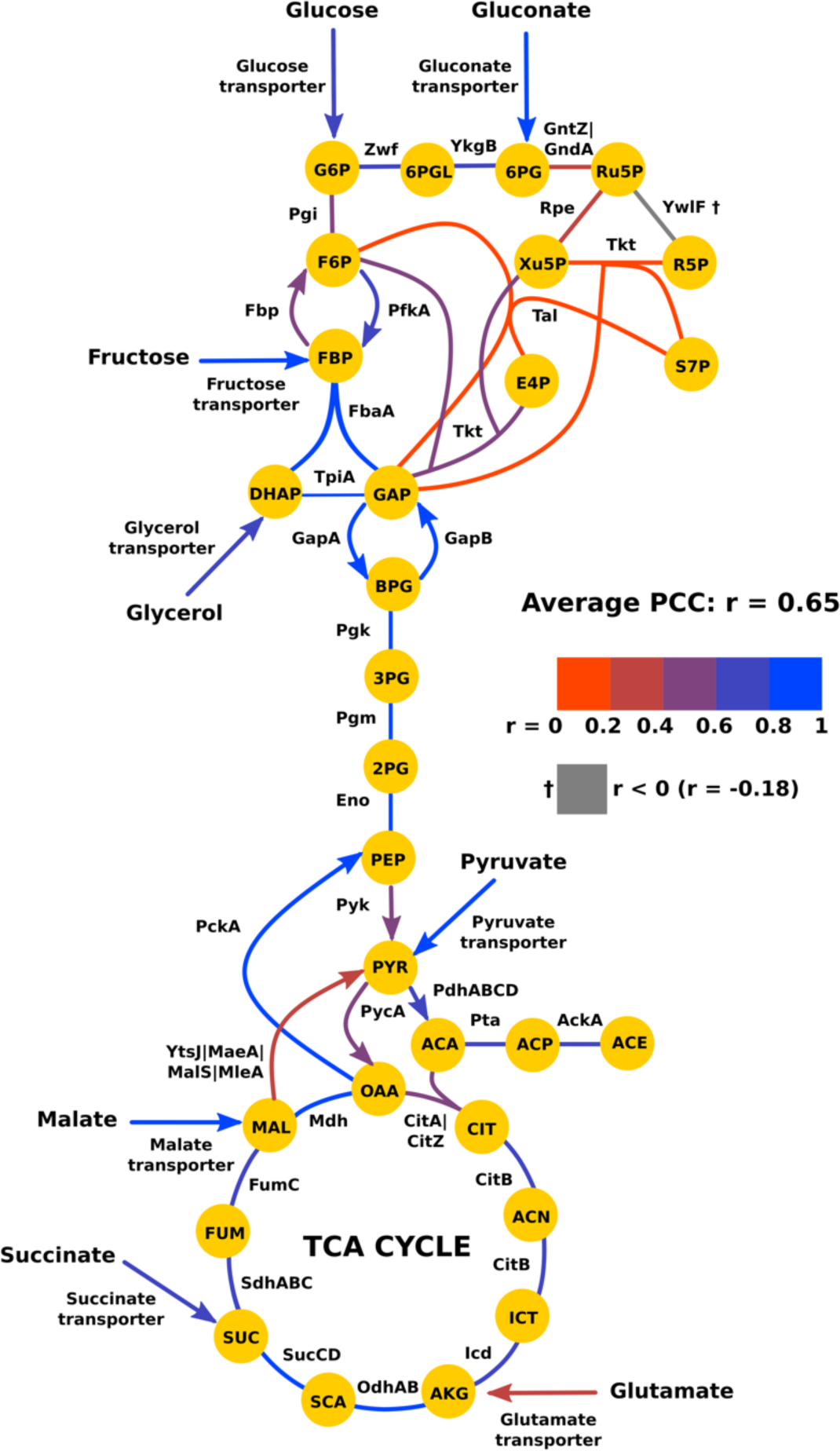
Color coded representation of the correlation between the flux prediction results and the gold standard (fluxes derived by ^13^C labeling experiments of Chubokov *et al* ^(26)^). The value r =1 represents the highest positive correlation (blue) between the flux prediction results and the gold standard, while r = 0 shows no correlation (red).

Next, we compared the distinct flux behavior of glycolysis and gluconeogenesis resulting from the investigated eight different growth conditions. *B. subtilis* utilizes two glyceraldehyde-3-phosphate dehydrogenases operating in opposite functions, i.e. glycolytic NAD-dependent GapA and gluconeogenic NADP-dependent GapB ^(38)^. Glucose and malate are the two main carbon sources for *B. subtilis* ^(39, 40)^. As expected, we observed glycolysis in our model, when *B. subtilis* was fed with glucose, i.e. a flux from glucose to the TCA cycle, through GapA (12.20 mmol h^-1^ gcdw^-1^). In turn, the opposite direction was observed when *B. subtilis* was fed with malate utilizing GapB (3.51 mmol h^-1^ gcdw^-1^). The fluxes derived from our model of both conditions are illustrated in Fig.4. For the other six conditions, in five out of six conditions, GapA and GapB were correctly predicted to be active based on the type of carbon sources (leading to glycolytic or gluconeogenic metabolism). Nevertheless, for the succinate/glutamate combined condition, the model showed a flux in GapA instead of GapB which was incorrect. This might be due to the fact that the second identified carbon source was predicted incorrectly (Table 2 and Fig.2). All predicted fluxes are provided in Table S7.

In summary, the model reflected the fluxes of glycolysis and TCA cycle quite well, but considerably less precisely for PPP.

**Figure 4.**
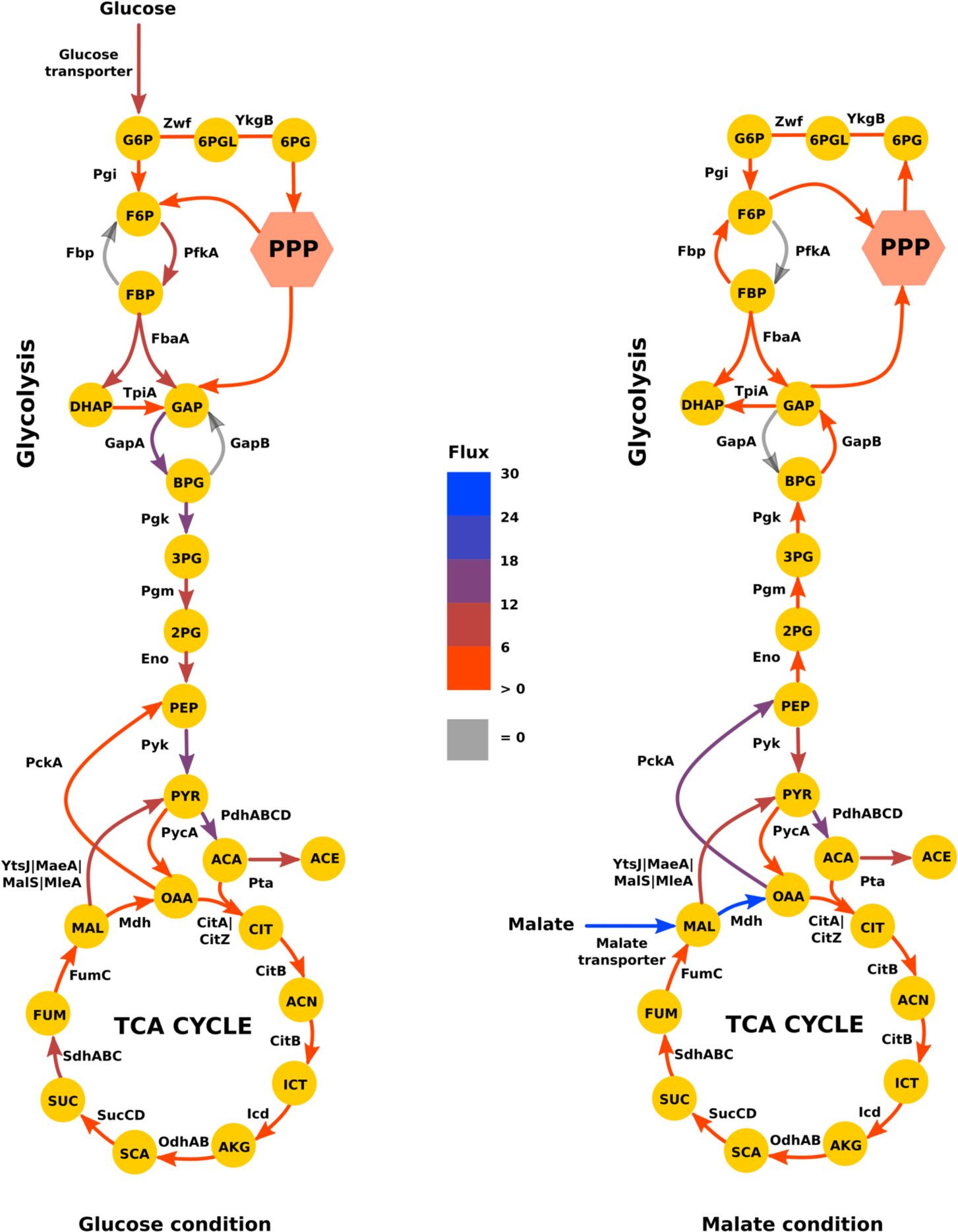
Exemplarily, the flux predictions from a glycolytic (glucose) and gluconeogenetic (malate) carbon source is shown. The level of the predicted fluxes is shown in mmol h^-1^ gcdw^-1^. For GapA, there is the predicted flux at the glucose condition (left), while at the malate condition (right) it is predicted to be no flux. In turn, for GapB, there is only the predicted flux for the malate condition.

### Gene expression profile based flux balance model identifies the carbon sources in an independent dataset of dynamic conditions

So far, we showed how we trained and tested our model utilizing gene expression data to predict carbon sources for *B. subtilis* in eight different steady-state conditions. However, in a natural environment, the bacteria may need to switch from one carbon source to another. Particularly, glucose and malate are preferred carbon sources for which a switch may occur ^(39, 40)^. We applied our approach to a publicly available time-series dataset consisting of two nutritional shifts, i.e. from glucose to glucose plus malate, and from malate to glucose plus malate. In these shifts, *B. subtilis* was grown on a single substrate, and the other substrate was added. Transcription profiles and metabolic flux data from ^13^C tracer analysis had been generated within the original study ^(24)^. For both time series experiments, quite close meshed data was available (before, and 5 min, 10 min, 15 min, 25 min, 45 min, 60 min, and 90 min after adding the other carbon source). We applied the model we have trained on the first dataset using the same parameter settings. We investigated the predictions of the major carbon sources for the two conditions before adding the other carbon source (only glucose, only malate), and for two conditions after the shift, i.e. 90 min after adding malate to glucose (glucose plus malate), 90 min after adding glucose to malate (malate plus glucose). They were considered most likely to be in a steady-state condition. Similar to the first dataset, we compared our predicted results with the available ^13^C metabolic flux data. Three out of four conditions (only glucose, only malate, and malate plus glucose) were predicted correctly. However, glucose plus malate was incorrectly predicted (Table 3A). We observed that there was a comparable amount of flux from fructose uptake during the shift from glucose to glucose plus malate. Going back to the carbon source predictions from the first dataset, we observed that the transporters at glucose condition and fructose conditions were quite similar (Fig.2). It is known that glucose and fructose transports share common coding genes (ptsI, ptsH) for enzyme I and HPr protein in sugar phosphotransferase system (PTS) ^(41)^. Thus, we reasoned the wrong prediction by assuming that the model could not distinguish between these two carbon sources in the glucose plus malate condition. To investigate this, we combined the uptake predictions from the glucose and fructose transporters and reassessed the comparison between the prediction results and ^13^C metabolic flux data. Indeed, after combining uptake predictions from glucose and fructose transporters (hexose), all four conditions (only glucose, glucose plus malate, only malate, and malate plus glucose) were predicted correctly (Table 3B).

**Table 3.** Prediction of the carbon source for the time series of the nutritional shift

**A**
| Transporter | Condition |  |  |  |
| --- | --- | --- | --- | --- |
|  | Glucose | GM<sup>a</sup> | Malate | MG<sup>a</sup> |
| Glucose | 1 | 1 | 2 | 1 |
| Malate | 2 | 2 | 1 | 2 |

**B**
| Transporter | Condition |  |  |  |
| --- | --- | --- | --- | --- |
|  | Glucose | GM<sup>a</sup> | Malate | MG<sup>a</sup> |
| Hexose | 1 | 2 | 2 | 1 |
| Malate | 2 | 1 | 1 | 2 |

<sup>a</sup> GM = Glucose to glucose plus malate at 90 min, MG = Malate to glucose plus malate at 90 min

Next, we tested our model in predicting the time-series behavior of the shifts. To obtain predictions in the dynamics of the shifts, we correlated the predicted fluxes of the malate and hexose transporters with the fluxes from the gold standard across all time points (Fig.S4). For the shift from malate to glucose plus malate, we obtained good correlations for both transporters, i.e. r = 0.80 for the hexose transporter, and r = 0.84 for the malate transporter indicating the correct behavior of the shift. For the shift from glucose to glucose plus malate, we observed a good correlation for the malate transporter (r = 0.83) but found a lower correlation for the hexose transporter (r = 0.32). The flux predictions from both shifts are provided in Table S11 and Table S12. In summary, the model performed well in predicting the carbon sources of the unseen dataset, when considering combining the predictions from the fructose and glucose transporters under one of the investigated conditions.

## DISCUSSION

We established a novel method to integrate gene expression profiles into the metabolic network. During implementing the method, we observed that the performance strongly depended on the clearance of thermodynamically infeasible loops (TILs) and an appropriate estimation of the lower and upper bounds of each reaction in the objective function. Hence, a major concern of our developments was to address these issues. We came up with a novel means for reducing the upper and lower bounds by iteratively reducing the flux ranges of each considered reaction. With this, the flux prediction results improved. Furthermore, we recognized that using an existing, well established means (ll-COBRA) to reduce TILs was very powerful, but, for our purposes quite slow. We addressed this issue and introduced our new method RED-TIL as an alternative method. While ll-COBRA formulates one large problem and searches for an optimal solution in a predefined-thermodynamic feasible region, RED-TIL splits the problem into smaller problems by detecting a TIL in the optimal solution, excluding it from the solution space and re-optimizing the solution iteratively until no TIL above a certain threshold is detected. We observed that both methods provided not exactly the same, but very similar and comparable results. More importantly, on average, RED-TIL removed TILs three times faster than ll-COBRA. To note, depending on the complexity of the problems, the speed-up varied between two to six times.

Using these improved means, we set-up an FBA model basing on a linear regression model between gene expression and predicted fluxes. Our main aim was to predict the main carbon source for the model organism *B. subtilis* and gauged the method with two independent datasets. As shown in our study, we succeeded in identifying the correct carbon sources for the different conditions based on the according gene expression profiles. Moreover, for many reactions in substrate uptakes, glycolysis, and TCA cycle, the flux prediction results correlated well with ^13^C metabolic flux data.

Besides being an alternative method to investigate metabolism, our approach is generalizable, flexible and versatile. It is unbound to any specific organism if for the organism of interest, there exists a well established metabolic network model and for the carbon source conditions under study, gene expression profiles are available. Our study supports the idea that FBA analysis based on gene expression profiles and following a concise goal like predicting the carbon sources, may serve as an alternative to ^13^C tracer analysis. In more complex situations where ^13^C labeling experiments are challenging to achieve, it may be a suitable means. Since our approach adjusts flux levels continuously to gene transcript levels, it circumvents defining thresholds to e.g. set genes as “expressed” or “non-expressed” as in Integrative Metabolic Analysis Tool (iMAT) ^(42)^ or in a Boolean logic setting like Gene Inactivity Moderated by Metabolism and Expression (GIMME) ^(43)^.

The method worked well for all except one condition. We discuss now this exception. Predicting the carbon source for the endpoint of the glucose to glucose plus malate shift (glucose plus malate) in the time-series dataset from Buescher *et al*, we could not predict glucose as the correct carbon source. When we studied the expression profiles of the according glucose transporters of the PTS operon and compared them to the respective ^13^C metabolic flux data from the glucose to glucose plus malate shift, the gene expression profile did not correlate with the according fluxes indicating different regulation mechanisms (Table S13). This observation was unusual, for most of our investigated transporters and carbon sources, we observed a good correlation (r > 0.60, see Fig.3). In addition, we observed a cross talk between glucose and fructose transporters in the first dataset (of Nicolas *et al*, eight carbon sources at steady state conditions) and circumvented this issue by regarding a representative of the combination of glucose and fructose transporters. This resulted in correct carbon source predictions leaving the prediction of the exact hexose indistinct. Notably, in the original publication, Buescher *et al* discussed that the adaptation induced by malate (glucose to glucose plus malate shift) does not primarily base on transcriptional regulation. Buescher *et al* performed multi-omics analysis using time lapse profiles from promoter activity, mRNA abundance, and protein abundance to identify post-transcriptional events ^(24)^. After correlating the gene expression level with the protein level, they observed high positive correlations in gene-protein pairs related to glycolysis such as phosphoglycerate mutase (r = 0.96), PTS glucose transporter (r = 0.88) and glyceraldehyde 3-phosphate dehydrogenase (r = 0.96) for the other shift, i.e. malate to glucose plus malate. However, they could not find correlations in gene-protein pairs related to glycolysis in the glucose to glucose plus malate shift. From this, they concluded that the discrepancy may be due to the initiation of translation during the glucose to glucose plus malate shift, and the shift was dominantly controlled by post-transcriptional mechanisms (in contrast to the malate to glucose plus malate shift). This is reasonable, as the benefit of glucose metabolism are very high for the bacteria, and it may be more beneficial for it to keep proteins for glycolysis constitutively expressed under the malate condition. We would add on this that taking the fructose transporters into account may compensate, to some extend, for this discrepancy. This sets a good example of the limitation of the approach. The method relies on gene expression profiles to predict the metabolic flux. It requires an in-depth investigation of the biological background beforehand to avoid studying mechanisms, which may rather depend on more direct regulation mechanisms and may be less dependent on transcriptional regulation.

## CONCLUSION

We have introduced a novel computational approach integrating gene expression profiles into a metabolic network including new methods to reduce flux ranges and remove TILs. With this, we could well predict the carbon sources of *B. subtilis* grown in each of these carbon sources. Our approach is promising, simple and generalizable.

## AUTHOR CONTRIBUTIONS

K.T., F.H. and M.O. developed the approach. K.T. carried out the project. K.T. and R.K. wrote the manuscript. All authors have read and approved the manuscript.

## ACKNOWLEDGEMENTS

We thank our research group members for helpful discussion and general support. We also thank Stefan Schuster for providing us useful suggestions on the project.

This work was supported by the German Federal Ministry of Education and Research (BMBF) within the project Center for Sepsis Control and Care (CSCC, 01EO1002 and 01EO1502), the Deutsche Forschungsgemeinschaft (https://www.dfg.de/) within the project KO 3678/5-1 and the Deutscher Akademischer Austauschdienst (DAAD). The funders had no role in study design, data collection and analysis, decision to publish or preparation of the manuscript.

The authors have declared that no competing interests exist.

## REFERENCES

1. McNeil JC, Vallejo JG, Kok EY, Sommer LM, Hulten KG, Kaplan SL. Clinical and Microbiologic Variables Predictive of Orthopedic Complications Following S. aureus Acute Hematogenous Osteoarticular Infections in Children. Clin Infect Dis. 2019.

2. Schmitt SK. Osteomyelitis. Infect Dis Clin North Am. 2017;31(2):325–38.

3. Zimmerli W, Sendi P. Orthopaedic biofilm infections. APMIS. 2017;125(4):353–64.

4. Kavanagh N, Ryan EJ, Widaa A, Sexton G, Fennell J, O’Rourke S, et al. Staphylococcal Osteomyelitis: Disease Progression, Treatment Challenges, and Future Directions. Clin Microbiol Rev. 2018;31(2).

5. Geraci J, Neubauer S, Pollath C, Hansen U, Rizzo F, Krafft C, et al. The Staphylococcus aureus extracellular matrix protein (Emp) has a fibrous structure and binds to different extracellular matrices. Sci Rep. 2017;7(1):13665.

6. Junka A, Szymczyk P, Ziolkowski G, Karuga-Kuzniewska E, Smutnicka D, Bil-Lula I, et al. Bad to the Bone: On In Vitro and Ex Vivo Microbial Biofilm Ability to Directly Destroy Colonized Bone Surfaces without Participation of Host Immunity or Osteoclastogenesis. PLoS One. 2017;12(1):e0169565.

7. Bouras D, Doudoulakakis A, Tsolia M, Vaki I, Giormezis N, Petropoulou N, et al. Staphylococcus aureus osteoarticular infections in children: an 8-year review of molecular microbiology, antibiotic resistance and clinical characteristics. J Med Microbiol. 2018;67(12):1753–60.

8. McNeil JC, Kaplan SL, Vallejo JG. The Influence of the Route of Antibiotic Administration, Methicillin Susceptibility, Vancomycin Duration and Serum Trough Concentration on Outcomes of Pediatric Staphylococcus aureus Bacteremic Osteoarticular Infection. Pediatr Infect Dis J. 2017;36(6):572–7.

9. Lewis NE, Schramm G, Bordbar A, Schellenberger J, Andersen MP, Cheng JK, et al. Large-scale in silico modeling of metabolic interactions between cell types in the human brain. Nat Biotechnol. 2010;28(12):1279–85.

10. Orth JD, Thiele I, Palsson BO. What is flux balance analysis? Nat Biotechnol. 2010;28(3):245–8.

11. Sharma AK, Konig R. Metabolic network modeling approaches for investigating the “hungry cancer”. Semin Cancer Biol. 2013;23(4):227–34.

12. Bideaux C, Montheard J, Cameleyre X, Molina-Jouve C, Alfenore S. Metabolic flux analysis model for optimizing xylose conversion into ethanol by the natural C5-fermenting yeast Candida shehatae. Appl Microbiol Biotechnol. 2016;100(3):1489–99.

13. Chiewchankaset P, Siriwat W, Suksangpanomrung M, Boonseng O, Meechai A, Tanticharoen M, et al. Understanding carbon utilization routes between high and low starch-producing cultivars of cassava through Flux Balance Analysis. Sci Rep. 2019;9(1):2964.

14. Dang L, Liu J, Wang C, Liu H, Wen J. Enhancement of rapamycin production by metabolic engineering in Streptomyces hygroscopicus based on genome-scale metabolic model. J Ind Microbiol Biotechnol. 2017;44(2):259–70.

15. Kavscek M, Bhutada G, Madl T, Natter K. Optimization of lipid production with a genome-scale model of Yarrowia lipolytica. BMC Syst Biol. 2015;9:72.

16. Chenard T, Guenard F, Vohl MC, Carpentier A, Tchernof A, Najmanovich RJ. Remodeling adipose tissue through in silico modulation of fat storage for the prevention of type 2 diabetes. BMC Syst Biol. 2017;11(1):60.

17. Gatto F, Miess H, Schulze A, Nielsen J. Flux balance analysis predicts essential genes in clear cell renal cell carcinoma metabolism. Sci Rep. 2015;5:10738.

18. Shan M, Dai D, Vudem A, Varner JD, Stroock AD. Multi-scale computational study of the Warburg effect, reverse Warburg effect and glutamine addiction in solid tumors. PLoS Comput Biol. 2018;14(12):e1006584.

19. Sung J, Kim S, Cabatbat JJT, Jang S, Jin YS, Jung GY, et al. Global metabolic interaction network of the human gut microbiota for context-specific community-scale analysis. Nat Commun. 2017;8:15393.

20. Antoniewicz MR. Methods and advances in metabolic flux analysis: a mini-review. J Ind Microbiol Biotechnol. 2015;42(3):317–25.

21. Lowe R, Shirley N, Bleackley M, Dolan S, Shafee T. Transcriptomics technologies. PLoS Comput Biol. 2017;13(5):e1005457.

22. Uygun S, Peng C, Lehti-Shiu MD, Last RL, Shiu SH. Utility and Limitations of Using Gene Expression Data to Identify Functional Associations. PLoS Comput Biol. 2016;12(12):e1005244.

23. van den Esker MH, Koets AP. Application of Transcriptomics to Enhance Early Diagnostics of Mycobacterial Infections, with an Emphasis on Mycobacterium avium ssp. paratuberculosis. Vet Sci. 2019;6(3).

24. Buescher JM, Liebermeister W, Jules M, Uhr M, Muntel J, Botella E, et al. Global network reorganization during dynamic adaptations of Bacillus subtilis metabolism. Science. 2012;335(6072):1099–103.

25. Nicolas P, Mader U, Dervyn E, Rochat T, Leduc A, Pigeonneau N, et al. Condition-dependent transcriptome reveals high-level regulatory architecture in Bacillus subtilis. Science. 2012;335(6072):1103–6.

26. Chubukov V, Uhr M, Le Chat L, Kleijn RJ, Jules M, Link H, et al. Transcriptional regulation is insufficient to explain substrate-induced flux changes in Bacillus subtilis. Mol Syst Biol. 2013;9:709.

27. Mudunuri U, Che A, Yi M, Stephens RM. bioDBnet: the biological database network. Bioinformatics. 2009;25(4):555–6.

28. King ZA, Lu J, Drager A, Miller P, Federowicz S, Lerman JA, et al. BiGG Models: A platform for integrating, standardizing and sharing genome-scale models. Nucleic Acids Res. 2016;44(D1):D515–22.

29. UniProt Consortium T. UniProt: the universal protein knowledgebase. Nucleic Acids Res. 2018;46(5):2699.

30. Kanehisa M, Furumichi M, Tanabe M, Sato Y, Morishima K. KEGG: new perspectives on genomes, pathways, diseases and drugs. Nucleic Acids Res. 2017;45(D1):D353–D61.

31. Kanehisa M, Goto S. KEGG: kyoto encyclopedia of genes and genomes. Nucleic Acids Res. 2000;28(1):27–30.

32. Kanehisa M, Sato Y, Kawashima M, Furumichi M, Tanabe M. KEGG as a reference resource for gene and protein annotation. Nucleic Acids Res. 2016;44(D1):D457–62.

33. van den Esker MH, Kovacs AT, Kuipers OP. YsbA and LytST are essential for pyruvate utilization in Bacillus subtilis. Environ Microbiol. 2017;19(1):83–94.

34. Benjamini Y, Hochberg Y. Controlling the False Discovery Rate: A Practical and Powerful Approach to Multiple Testing. Journal of the Royal Statistical Society Series B (Methodological). 1995;57(1):289–300.

35. Mahadevan R, Schilling CH. The effects of alternate optimal solutions in constraint-based genome-scale metabolic models. Metab Eng. 2003;5(4):264–76.

36. Price ND, Famili I, Beard DA, Palsson BO. Extreme pathways and Kirchhoff’s second law. Biophys J. 2002;83(5):2879–82.

37. Schellenberger J, Lewis NE, Palsson BO. Elimination of thermodynamically infeasible loops in steady-state metabolic models. Biophys J. 2011;100(3):544–53.

38. Fillinger S, Boschi-Muller S, Azza S, Dervyn E, Branlant G, Aymerich S. Two glyceraldehyde-3-phosphate dehydrogenases with opposite physiological roles in a nonphotosynthetic bacterium. J Biol Chem. 2000;275(19):14031–7.

39. Kleijn RJ, Buescher JM, Le Chat L, Jules M, Aymerich S, Sauer U. Metabolic fluxes during strong carbon catabolite repression by malate in Bacillus subtilis. J Biol Chem. 2010;285(3):1587–96.

40. Meyer FM, Stulke J. Malate metabolism in Bacillus subtilis: distinct roles for three classes of malate-oxidizing enzymes. FEMS Microbiol Lett. 2013;339(1):17–22.

41. Stulke J, Hillen W. Regulation of carbon catabolism in Bacillus species. Annu Rev Microbiol. 2000;54:849–80.

42. Zur H, Ruppin E, Shlomi T. iMAT: an integrative metabolic analysis tool. Bioinformatics. 2010;26(24):3140–2.

43. Becker SA, Palsson BO. Context-specific metabolic networks are consistent with experiments. PLoS Comput Biol. 2008;4(5):e1000082.

