## Supplementary Information for "Gene expression profiles based flux balance model to predict the carbon source for *Bacillus subtilis*"

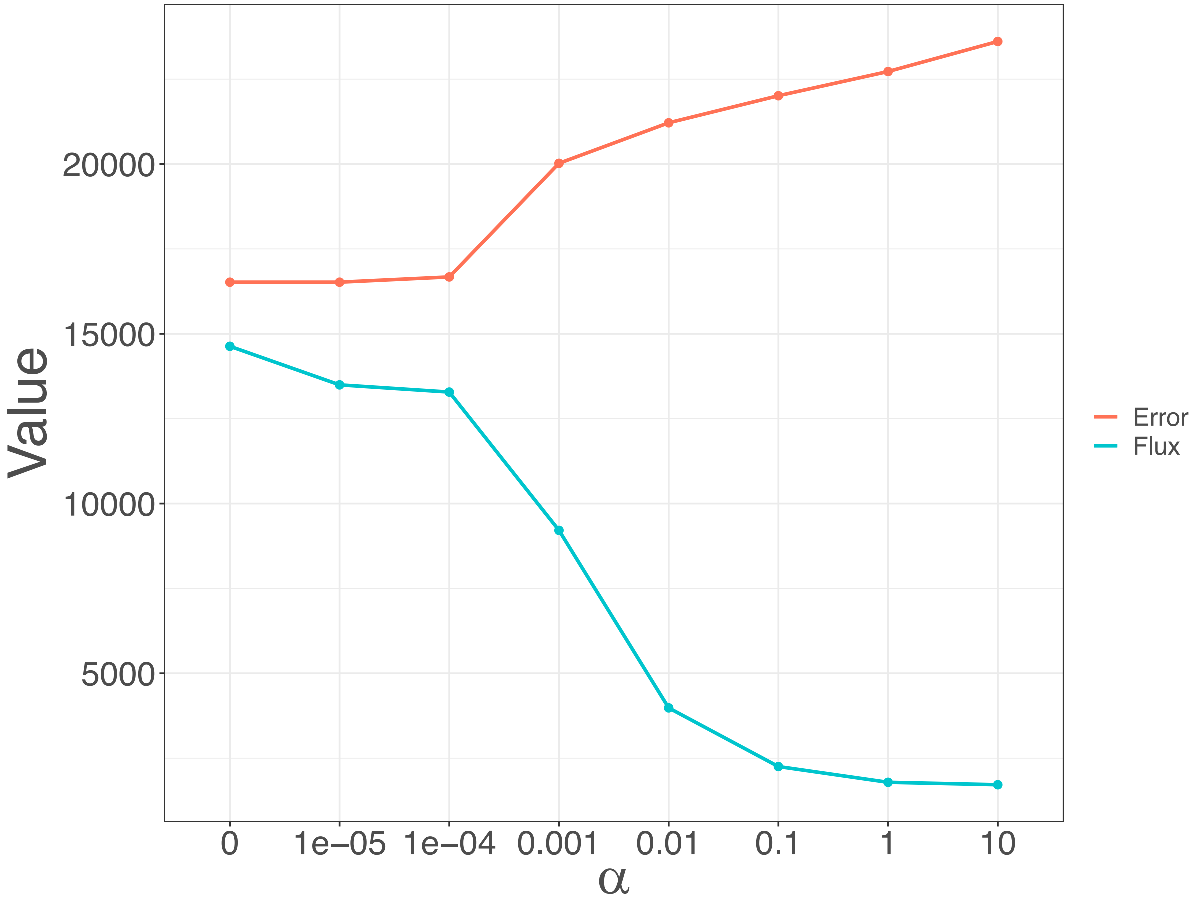


**Figure S1** Trade-off between the sum of flux values from reactions which are not core nor associated reactions and total model mapping error calculated from all eight conditions at different $\alpha$ values. At $\alpha = 0.01$, the sum of flux values drops dramatically while the total model mapping error only slightly increases.

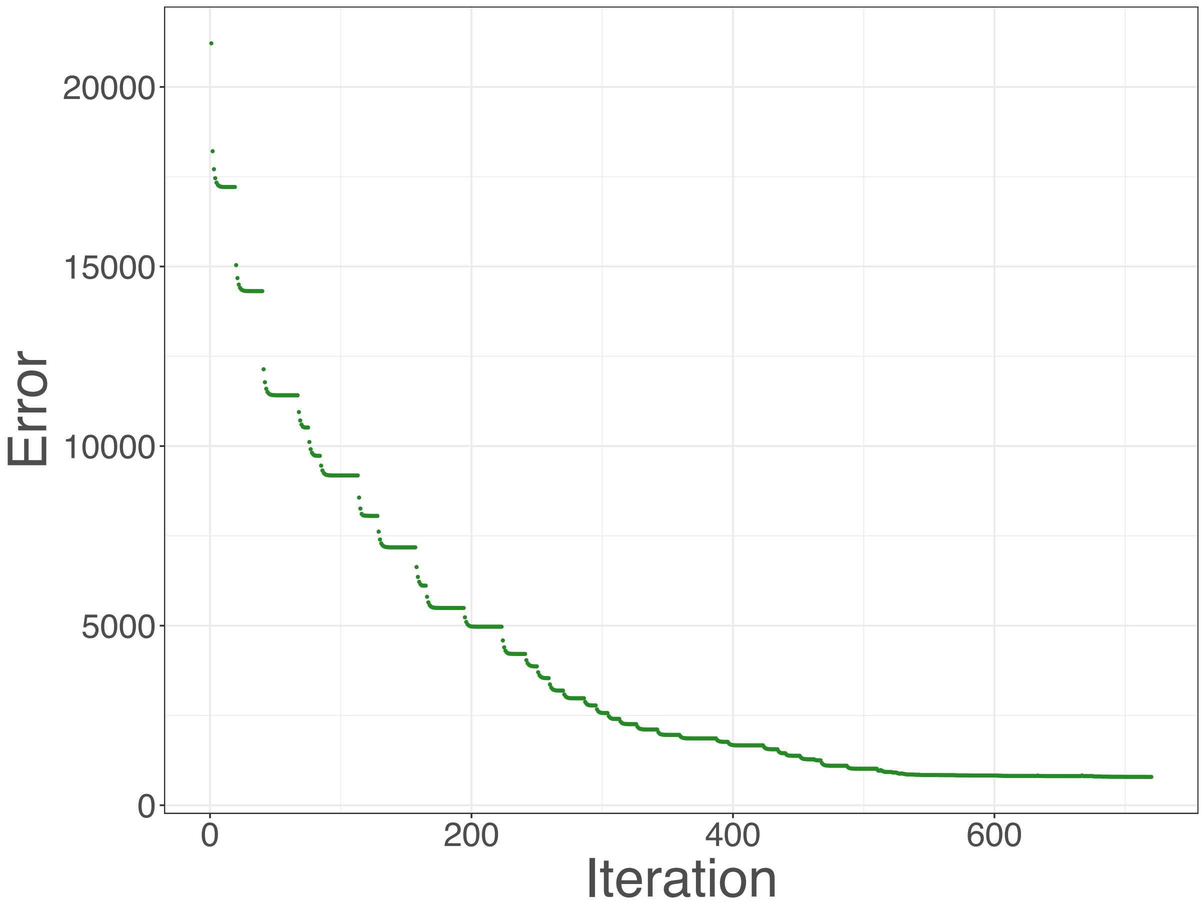


**Figure S2** The total model mapping error calculated from all eight conditions at $\alpha$ = 0.01 is shown with respect to the number of iterations of the search space reduction algorithm. As the algorithm proceeds near the end of the search space reduction list (>500 iterations), the total model mapping error does not further decrease.

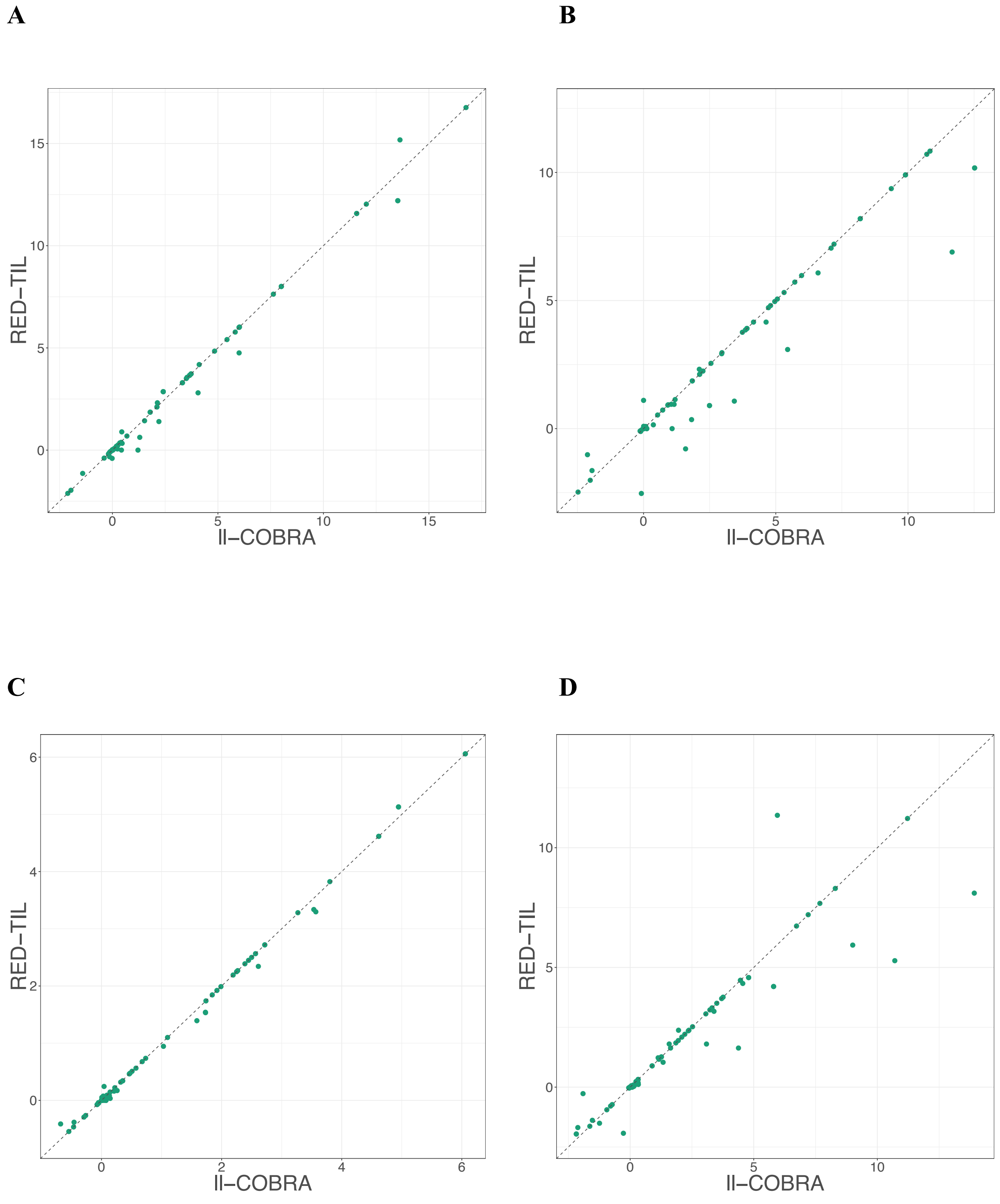
A
B
C
D
RED−TIL
Il−COBRA
0
5
10
15
−5

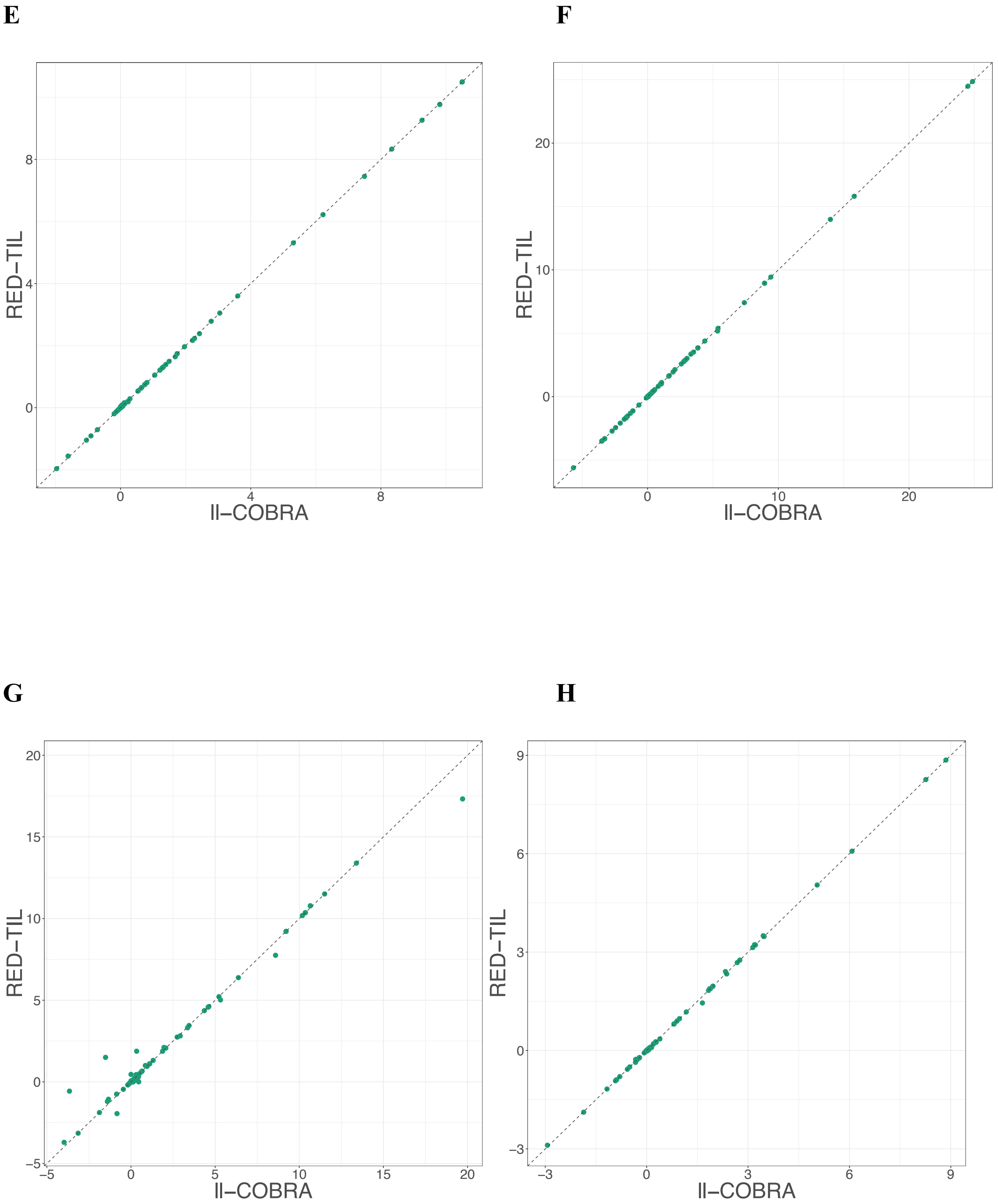


**Figure S3 (A-H)** Scatterplots comparing the predicted fluxes of the core and associated reactions (98 reactions) from ll-COBRA with the predicted fluxes from RED-TIL. The conditions are glucose (A), fructose (B), gluconate (C), glutamate/succinate (D), glycerol (E), malate (F), malate/glucose (G), and pyruvate (H).

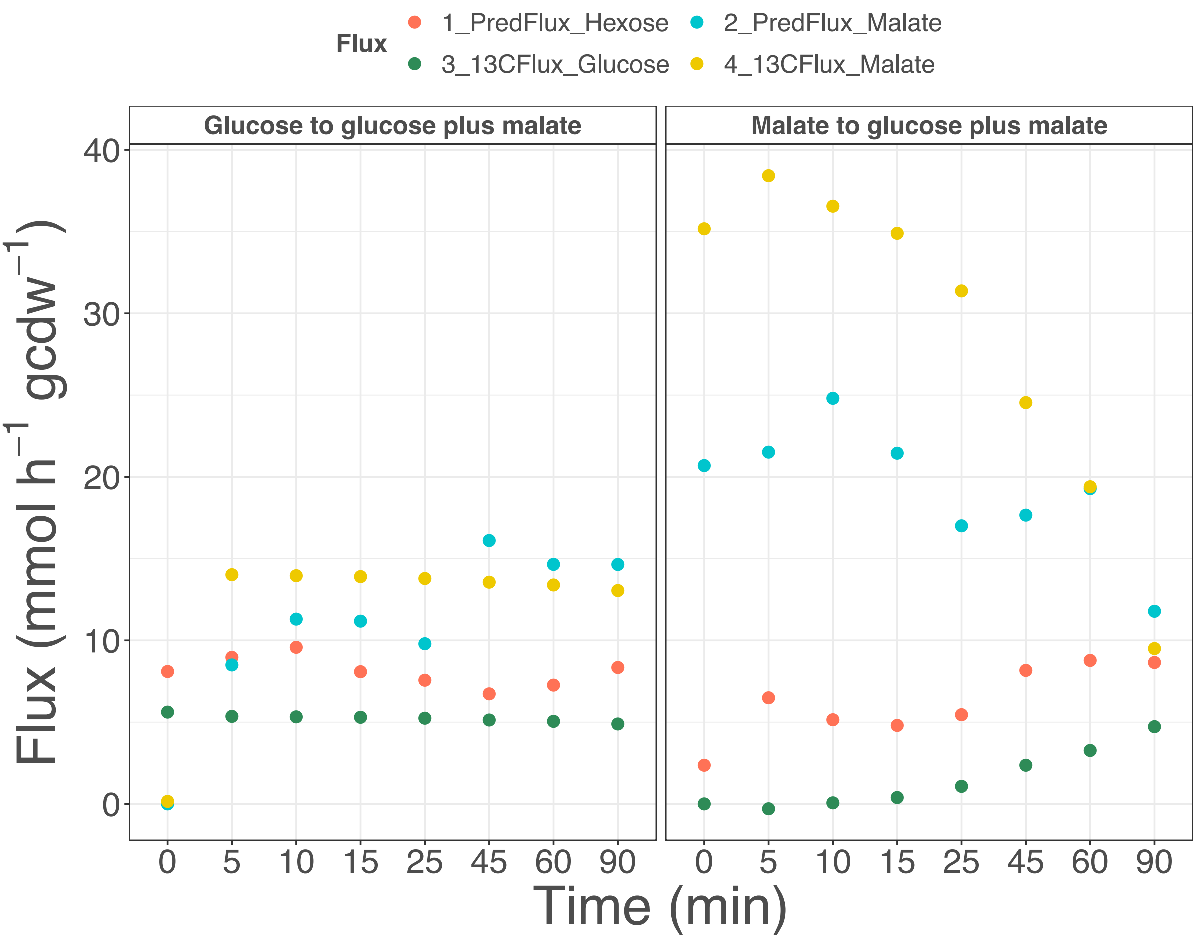


**Figure S4** Carbon source shifts between glucose and malate. Red dots (PredFlux_Hexose) represent the flux prediction results from the hexose transporters, while blue dots show the malate transporter (PredFlux_Malate). However, green and yellow dots represent the fluxes derived from $^{13}C$ tracing (from the original publication) (13CFlux_Glucose and 13CFlux_Malate, respectively). After adding the second substrate, the uptakes from glucose and malate transporter start to change.

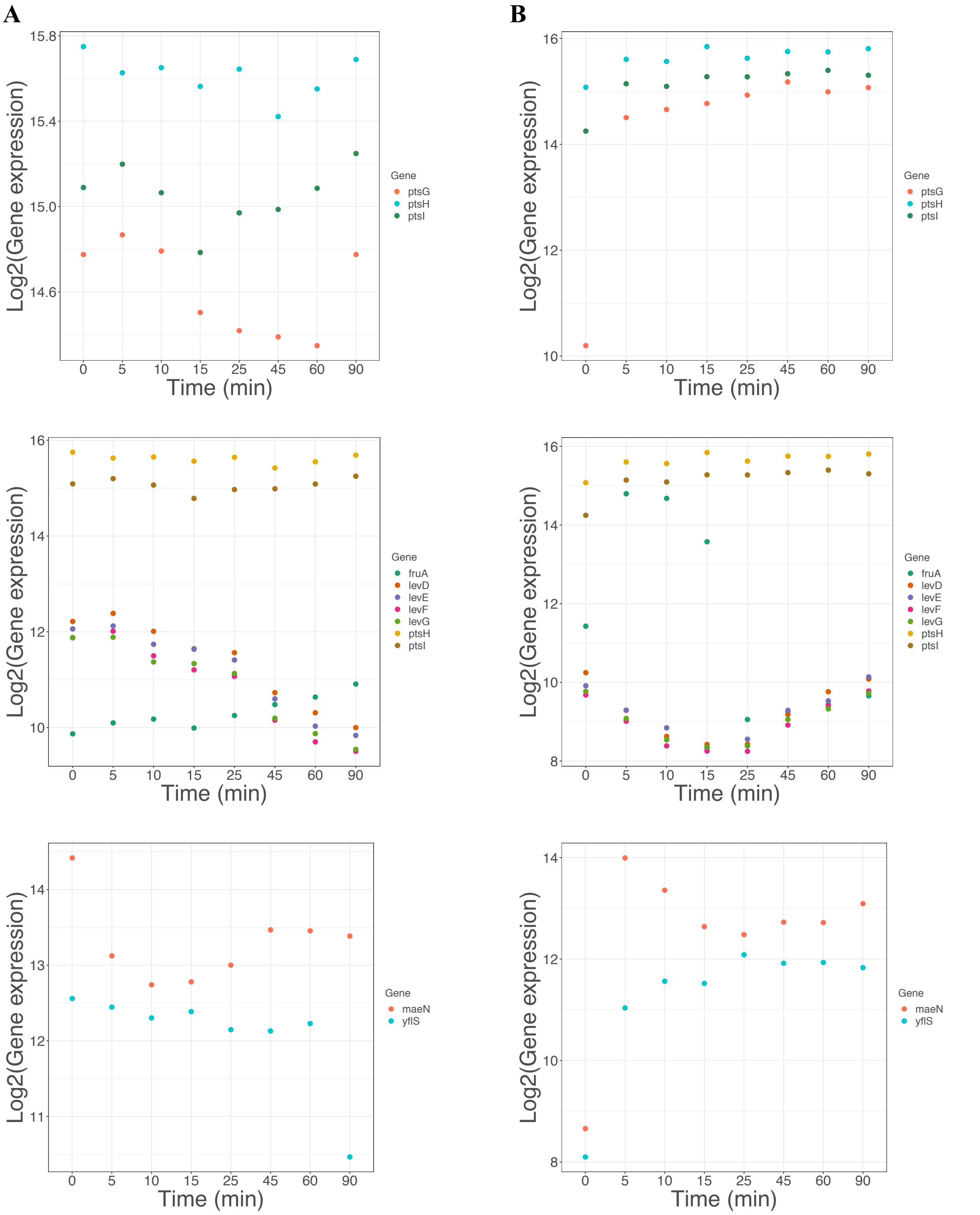


**Figure S5 (A and B)** Gene expression profiles of genes involved in glucose (ptsG, ptsH, ptsI), fructose (fruA, ptsH, ptsI, levD, levE, levF, levG) and malate (maeN, yflS) transporter reactions from glucose to glucose plus malate (A) and malate to glucose plus malate (B) for each time point.

**List of supplementary tables**

All supplementary tables (Table S1-S13) are provided in the supplementary table file (.xlsx).
